# Powerful solution for experimental cerebral malaria treatment: artesunate and tetramethylpyrazine

**DOI:** 10.1101/622688

**Authors:** Xiaohui Jiang, Lina Chen, Zhongyuan Zheng, Yuan Guo, Kai Li, Ying Chen, Xiaogang Weng, Ting Yang, Shuiqing Qu, Hui Liu, Yujie Li, Xiaoxin Zhu

## Abstract

**Background:** Cerebral malaria (CM) is a kind of serious neurological complication caused by the acute Plasmodium falciparum infection. About 300000 patients including children under 5 years old died from this disease every year. Even intravenous artesunate (Art) is employed as the most effictive drug in the treatment of CM, high incidence of death and neurological sequelae are still inevitable. Therefore, we assessed the combination of Art and tetramethylpyrazine (TMP), to treat experimental CM (ECM) in C57BL/6 mice infected with *Plasmodium berghei* ANKA (PbA). A non-biased whole brain quantitative proteomic analysis was also conducted to get some insight of the mechanism of the combinational treatment.

**Results:** Treatment of (ECM)-C57BL/6 mice with the combination of Art and TMP increased the survival, improved clinical signs and prevented neurological manifestations. These effects were related to reduction of parasitised red blood cells (pRBC) adhesion, sequestration, maintaining brain microvascular integrity, increasing nerve growth factor, neurotrophin levels, and alleviating hippocampal neuronal damage and astrocyte activation. The pharmacological effects of Art-TMP combination therapy were analyzed by ECM mice brain proteomic function enrichment. Based on an isobaric tag for relative and absolute quantitation (iTRAQ) fold-change of 1.2 (P-value < 0.05), 217 down-regulated and 177 up-regulated proteins were identified, presenting a significantly altered proteome profile of the combined Art and TMP group as compared to the group treated with Art or TMP alone. These results suggested that the Art-TMP combination could be used as a powerful solution for CM and its neurologic damage.

**Conclusions:** An effictive therapy for CM with low mortality rate and protect against ECM-induced neurocognitive impairment has been proposed through the combination of Art and TMP, which can provide an effective adjuvant treatment in the clinic. iTRAQ proteomics provide a resource for further mechanistic studies to examine the synergistic effects of Art and TMP and their potential to serve as an adjunctive treatment method and intervention targets.

**Author Summary:** Cerebral malaria (CM) is the most serious neurological complication caused by Plasmodium falciparum infection. Even after antimalarial treatment, severe neurological sequelae still exist. We used tetramethylpyrazine (TMP), the main ingredient of the traditional Chinese medicine Chuanxiong, and artesunate (Art) as a combination of drugs. We found that Art-TMP combination could improve the clinical symptoms of CM and protect the nervous system. At the same time, proteomics was used to analyze the protective mechanism of Art-TMP combination administration on ECM mice. This study suggests that the combination of Art and TMP may be used as an adjuvant therapy for clinical CM and iTRAQ proteomics provides resources for further study of Art-TMP combination and provides potential prognostic biomarkers for this therapeutic intervention.

## Introduction

Malaria is considered to be one of the world’s three major infectious diseases that resulted in the death of 435,000 people in 2017. According to the 2018 World Malaria Report, 61% of all deaths due to malaria occurred in children under the age of five years [1]. Cerebral malaria (CM) is one of the major complications of *Plasmodium falciparum* infection. It is associated with severe disturbances of the consciousness (deep coma) and respiratory distress or other neurological abnormalities [2]. The World Health Organization (WHO) has recommended intravenous Art as the first-line treatment for severe malaria. Moreover, the therapeutic effects of intravenous Art are superior to those of quinine [3, 4]. However, Art monotherapy is insufficient to control the mortality rate, owing to a lack of specific neuro- and vasculo-protective effects, leading to approximately 300,000 deaths each year. Furthermore, about 26% of individuals present neurological deficits, such as learning and memory deficits, and language disability despite being administered anti-malarial drugs [5, 6]. Moreover, intravenous administration is largely restricted to the high-risk areas of malaria in Africa, CM remains a dominant cause of mortality and neurodisability in the tropics. As a consequence, there is an urgent clinical need to develop more effective and robust treatments for CM, with an aim to improve the protective effects of anti-malarial drugs.

Literature regarding the cerebral processes involved in the pathogenesis of CM and those that undermine the recovery from the complication after anti-malarial drug therapy is scarce. However, there is an increasing consensus that protecting the host brain vascular system and neurons and modulating the host pro-inflammatory immune response to infection are plausible effective strategies to improve the success of the anti-malarial drug therapy of CM. Indeed, vascular obstruction and immunopathology are known to be generally associated with the life-threatening symptoms of CM, which may interrupt the recovery by activating endothelial cells, astrocytes, and microglial cells in the brain, thereby disrupting the blood–brain barrier integrity, disturbing and/or destroying neuronal signalling and causing nerve injuries [7, 8]. All these abnormalities have been observed in the brains of patients with CM. It is believed that a dysfunction of blood vessels in the brain is the primary cause of development of CM, which may prevent the re-establishment of brain homeostasis, leading to the failure of the anti-malarial drug treatment.

To improve the fatal outcome caused by CM, the need of the hour is to explore a novel, inexpensive adjunctive therapy that can be easily administered and can improve neurological sequelae. Although Chuanxiong is not a major therapeutic medicine for malaria, several reports on the prescription of Chuanxiong as a combination therapy for malaria exist in the classical literature, such as nasal plug of Chuanxiong for malaria in Mongolian medicine. Tetramethylpyrazine (TMP) is the primary active alkaloid component of the traditional Chinese medicine Chuanxiong. Its use to treat cerebrovascular diseases could be traced back to thousands of years ago; moreover, it is known to exert protective effects on various nervous system injuries [9, 10]. TMP has been demonstrated to increase cerebral blood flow, improve microcirculation, inhibit the production of pro-inflammatory factors and protect learning and memory functions [11, 12]. Recently, clinical studies have reported that TMP exerts beneficial effects on the nervous system that can promote functional recovery from nerve injury. Adjunctive therapy is defined as an additional treatment that modifies the pathological processes caused by malaria to improve its clinical outcomes and/or decline mortality, along with the prevention of long-term neurocognitive impairment [13]. Over the past decades, dozens of CM interventions for different pathways have been evaluated based on immunomodulation, neuroprotection, regulation of gaseous signalling molecules and improvement of endothelial dysfunction [14]. In fact, more than 17 clinical trials have so far examined 11 treatments; however, no method has been proven effective in treating CM in children [15, 16].

The present study aimed to find an adjunctive therapy to improve neurological symptoms and survival in an experimental cerebral malaria (ECM) model. C57BL/6 mice were infected with *Plasmodium berghei* ANKA (PbA) and treated daily with Art, TMP, and a combination of Art and TMP. Parasitaemia and clinical, histological, and immunological features of the disease were monitored. Subsequently, to elucidate possible targets of Art-TMP combination in treating ECM, we performed an extensive quantitative proteomic analysis to compare the brain proteome profiles of ECM mice treated with Art, TMP, and Art-TMP combination with those of ECM model control mice. Interestingly, certain brain proteins, such as Slit2, Tiam2, Syntenin, and Hemopexin, were found to be sequentially altered in mice treated with the Art-TMP combination as compared to the proteins in the ECM mice. Here, we present the first comprehensive brain proteomic analysis of ECM mice treated with different drug combinations.

## Materials and Methods

### Ethics Statement

All experiments were carried out to minimize the suffering of animals. The care of laboratory animal and the animal experimental operation have conforming to Beijing Administration Rule of Laboratory Animal. All animal experiments were approved by the Animal Experimental Ethics Review Committee of the Institute of Basic Research for Chinese Medicine, China Academy of Chinese Medical Sciences (license number: SYXK (Beijing) 2016-0021).

### Mice, parasitic infection, and disease assessment

Six-to eight-week-old male C57BL/6 mice weighing 14 to 16 g were purchased from the National Institutes for Food and Drug control (Beijing, China). *P. berghei* ANKA, originally obtained from Dr. Ya Zhao at the Fourth Military Medical University, was passaged *in vivo*. Experimental mice were infected with 1 × 10^6^ parasitised red blood cells (pRBC) via intraperitoneal (i.p.) injection (recorded as day 0 [d0] post infection [p.i.]). Parasitaemia was monitored every day using Giemsa-stained blood smears. When mice develop reduced responsiveness to external stimuli, ataxia, paralysis or coma and convulsions are considered as typical symptoms of ECM [17].

### Treatment and drug administration

Experimental mice were randomised into four groups: PbA-infected group (Infected); artesunate (Art)-treated group; tetramethylpyrazine (TMP)-treated group and Art-TMP combination (Art+TMP)-treated group. Mice were treated with intranasal administration (IN) of 13.00 mg·kg^−1^ artesunate in the Art group, with ligustrazine hydrochloride injection (10.40 mg·kg^−1^) in the TMP group, and with Art-TMP combination (23.40 mg·kg^−1^) in the Art+TMP group, daily, starting from d2 till d5 p.i..

### Basic indicator evaluation

To evaluate the effect of drugs on mice, survival rate (SR), body weight and body temperature were measured. Parasitaemia was monitored using the Giemsa-stained blood smears, and clinical symptoms of the diseased mice were evaluated using the rapid murine coma and behaviour scale (RMCBS) from d3 p.i. The RMCBS consists of 10 parameters (gait, balance, motor performance, body position, limb strength, touch escape, pinna reflex, toe pinch, aggression and grooming). Each parameter is scored 0 to 2, with a 0 score correlating with the lowest function and a 2 score correlating with the highest [18]. The lower the score, the worse the state of the mouse, thus indicating severity of the disease.

### The open field test

We used the open field test to observe the spontaneous activity characteristics of mice entering a new environment [19]. Here, we used normal mice of the same age as control. Briefly, mice were placed in four drums of an empty field activity test box (an area with a central radius of 7.5 cm in a barrel is considered as the central area). Mice were placed in the empty field and spontaneous activities of the animals were observed within 5 minutes and recorded by the software automatically.

### Y-maze spontaneous alternation test

Spontaneous alternation in the Y-maze was performed as previously described [20]. A computer-controlled infrared camera system was installed directly above the Y maze to track the location of mice. Animals were placed in a fixed position of the Y maze in turn and allowed to explore freely for 8 minutes; normal mice were also used as a control. The total number of entries into each arm was recorded during the experiment.

### Histopathology and immunohistochemistry

Six mice in each group were killed at 7 d p.i. Tissues were stained with haematoxylin and eosin (H&E) for detecting microvascular obstructions. Immunohistochemical staining was performed with specific polyclonal antibodies against intercellular adhesion molecule-1 (ICAM-1), vascular cell adhesion molecule-1 (VCAM-1), glial fibrillary acidic protein (GFAP) and neuron-specific nuclear protein (NeuN) to detect the protein expression. Vessels were counted in 20 randomly selected fields per mouse using the method described in the literature [21]. GFAP-immunopositive cells in the cerebral cortex and NeuN-immunopositive cells in the hippocampus were quantified in each mouse brain and counted in five areas per section. The averaged data were used to evaluate the infiltration of inflammatory cells into viable the neural tissue and to detect viability of neurons.

### Assessment of vessel integrity and patency by magnetic resonance imaging

Mice in four groups were imaged at d7 p.i. using the following protocol. Each mouse was anesthetised with 2,2,2-tribromoethanol (0.32 mg/kg^−1^). Vessel integrity and patency was scanned using TOF-2D-FLASH employing a magnetic resonance imaging [MRI] scanner (BioSpec70/20 USR; Bruker, Germany) with the following parameters: flip angle = 50 degrees, field of view = 20 × 20 cm^2^, acquisition matrix size = 256 × 256, slices = 0.5 mm, repetition time = 10 ms, echo time = 1.84 ms, number of excitations = 5, imaging time = 6 minutes 49 seconds.

### Cytokine antibody arrays

A panel of mouse cytokines was assessed by determining their relative levels of expression using the RayBio® Cytokine Antibody Arrays (RayBiotech; Guangzhou, China). In brief, 24 brain tissue samples were lysed and adjusted to a final concentration of 500 μg/mL. Next, the cytokine antibody array was applied according to the manufacturer’s protocol. Intensities of fluorescent signals of the microarray were measured using a laser microarray scanner (InnoScan 300 Microarray Scanner; Innopsys; France) at 532 nm and quantified using the RayBio® Analysis tool software.

### Measurement of cytokine by enzyme-linked immunosorbent assay

The levels of cytokines including brain-derived neurotrophic factor (BDNF), neurotrophic factor-3 (NT-3) and tumour necrosis factor (TNF-α) were determined in the mice brain using commercial enzyme-linked immunosorbent assay (ELISA) kits as per the manufacturer’s protocol. The absorbance value was measured at 450 nmusing the Thermo Mulitskan MK3 microplate. The concentration of each neurotrophic factor in every sample was calculated via a standard curve generated using recombinant cytokines. Data are represented as mean ± SEM from six animals in each group.

### iTRAQ-based quantitative proteomic analyses

Specific methods of iTRAQ-based quantitative proteomic analyses were conducted using a method similar to that described in the literature [22]. In brief, the sample was lysed[23] followed by homogenisation; proteins in the sample were run on a sodium dodecyl sulphate polyacrylamide gel electrophoresis. The run sample was subjected to filter-aided sample preparation digestion, iTRAQ labelling, peptide fractionation using strong cation exchange chromatography, followed by HPLC and LC-MS/MS and data analyses.

### Functional enrichment analysis

To further explore the impact of differentially expressed proteins and discover the internal relations between them, an enrichment analysis was performed. GO enrichment on three ontologies (biological processes, molecular functions, and cellular components) and the KEGG pathway enrichment were applied using Fisher’s exact test, considering the whole quantified protein annotations as the background dataset. Benjamini–Hochberg correction for multiple testing was further applied to adjust the derived P-values. Only functional categories and pathways with P-values under a threshold of 0.05 are considered significant. The protein–protein interaction information of the studied proteins was retrieved using the STRING software (http://string-db.org/).

### Immunoblot analysis

Proteomic results were confirmed by western blotting using the BIORAD system or simple western analysis using the Wes^TM^ protocol according to the manufacturer’s (Protein Simple Biosciences & Technology, USA) protocol. All primary antibodies were used under the same condition (1:1000 dilution). For western blot analysis, the lysed protein was denatured, followed by loading of the same amount of protein of each mouse. The sample was electrophoresed, transferred, blocked, and incubated with rabbit anti-mouse primary antibody overnight at 4°C. The sample was then incubated with a secondary antibody at room temperature for 2 hours, after which enhanced chemiluminescence (ECL) chromogen was added, the blot was scanned and analysed in a gel-imaging system with actin as the internal reference.

For simple western analysis by Wes^TM^, brain homogenate samples were prepared, and protein concentration was determined using the bicinchoninic acid kit. The brain homogenate samples were diluted to a final concentration of 1 μg/µL as required by Wes^TM^. Capillary electrophoresis immunoassay was performed using Wes-specific reagent, 12–230 kDa Wes separation module 8 × 25 capillary filter. Electrophoresis images were generated using the Protein Simple Compass software.

### Nasal administration toxicity study

Toxicity study of nasal administration (NA) was assessed by histopathology of the nasal mucosa. Nine healthy C57BL/6 mice were administered a placebo solution or a mixed solution of injectable Art and ligustrazine hydrochloride injection (20 mg·kg^−1^) nasally. Three control mice and three mice administered drugs were sacrificed 30 minutes later. The other three mice were sacrificed 4 days after the intranasal administration of a mixture of injectable Art and ligustrazine hydrochloride injection (20 mg·kg^−1^). The nasal septum mucosa was collected, and the blood and mucus were washed with saline. These were fixed in 4% paraformaldehyde solution, dehydrated with gradient ethanol followed by HE staining. The paraffin-embedded sections were observed for histopathological changes in the nasal mucosa.

### Statistical analysis

All data are presented as mean of each group ± SEM. Data were analysed by one-way analysis of variance (ANOVA) and non-parametric tests, followed by Dunn’s multiple comparison or Tukey’s multiple comparison test. GraphPad prism version 5.0 (GraphPad) was used for charting and statistical analysis. P-values less than 0.05 are considered statistically significant.

## Results

### Effect of Art-TMP combination on mortality, parasitemia, and malaria parasite morphology in C57BL/6 mice infected with PbA

C57BL/6 mice infected with PbA ECM developed neurological signs on d6 p.i. that resulted in death between d8 and 12 p.i. (Fig. 1A, B). In contrast, only approximately 30% mice died in the Art group; deaths were observed from d10 p.i. The TMP treatment group did not improve the survival or reduce parasitemia and death occurred on d9 to 12 p.i.. Art + TMP treatment significantly improved the outcome in ECM mice; none of the mice in this group died and parasitaemia was reported only in 10.12% mice on d12 p.i. In addition, malaria parasites could still be observed in the Art group. A decrease in late trophozoites was observed in the blood smears of the Art + TMP group as compared to the blood smears of the Infected group and TMP alone treatment group on d7 p.i. (Fig. 1C).

**Figure 1.**
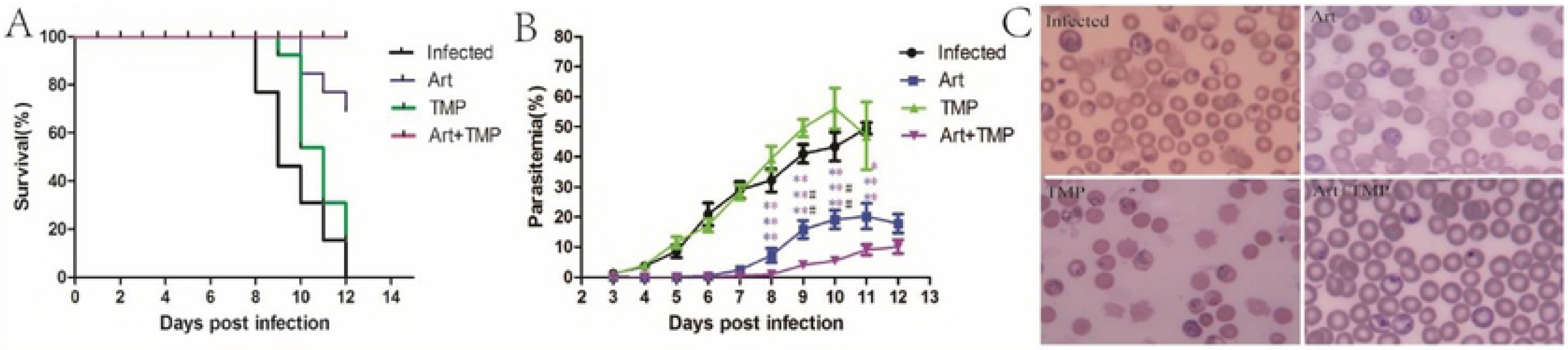
Effect of Art-TMP combination on mortality, parasitaemia and malaria parasite morphology in mice infected with PbA-ECM. Blood was collected from the tail of mice and stained with Giemsa for analysis of parasitaemia. Daily survival was also recorded. All ECM mice were treated with a single dose of Art at 13.00 mg·kg-1, a single dose of TMP at 10.40 mg·kg-1 and a dose of Art-TMP combination at 23.40 mg·kg-1 from d2 to d5 p.i..(A) Art-TMP combination significantly prolonged the survival of ECM mice. (B) Effect of Art and Art + TMP on parasitaemia in PbA-infected mice. (C) Giemsa-stained blood smears of PbA-infected mice on day 7 p.i. (n = 13 for each group). Data are expressed as mean ± SEM in each group. The parasitaemia was analysed by one-way ANOVA. Statistical differences associated with the infected group were marked according to the colour of the symbol of each group. ANOVA, analysis of variance; Art, artesunate; ECM, experimental cerebral malaria; PbA, Plasmodium berghei ANKA; SEM, standard error of mean; TMP, tetramethylpyrazine.

### Art-TMP combination reduces clinical symptoms of ECM

We next utilized rapid murine coma and behaviour scale (RMCBS) to assess ECM manifestations after drug therapy. We found that ECM mice developed a score less than 4 points, consistent with their symptoms of reduced exploratory behaviour, decreased reflex, self-preservation, and finally coma and epilepsy (Fig. 2A). The score in the Art-treated mice was significantly higher than that in the ECM mice in later stages of infection. Particularly, mice in the Art + TMP group presented distinctly higher RMCBS values as compared to ECM mice (Fig. 2A). Results also showed significant differences between the Art + TMP and the Art groups at 9 to 11 days p.i. The value was higher than 12 points in the Art + TMP group on d12 p.i. (Fig. 2A).

**Figure 2.**
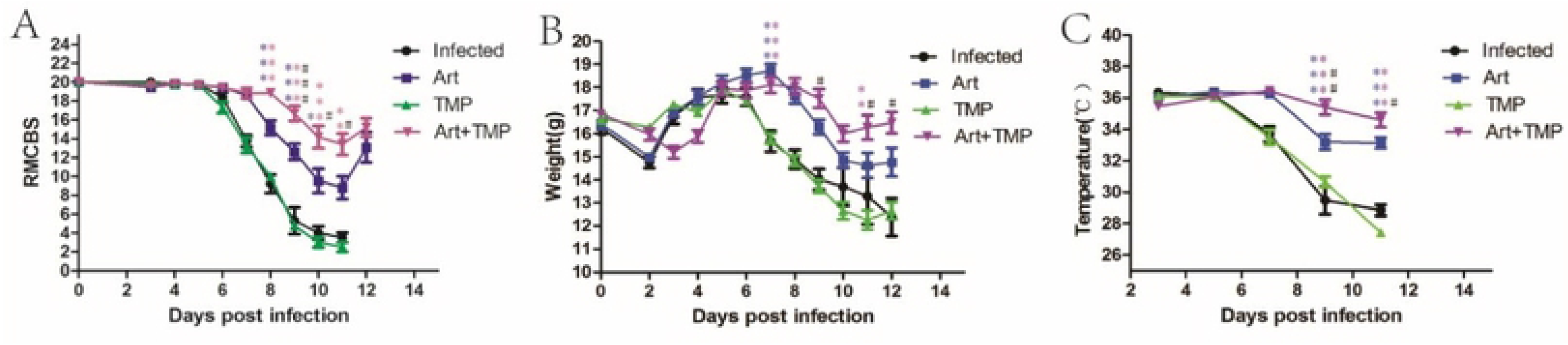
Art-TMP combination treatment can provide relief from clinical symptoms in ECM mice and improve weight and body temperature of ECM mice (n = 13). The weight of each mouse was counted daily from d2 p.i., RMCBS score evaluation was performed from d3 p.i. and body temperature was measured every other day. All ECM mice were treated with drugs as described above. (A) RMCBS score, (B) body weight and (C) body temperature of each group are shown. Data are expressed as mean ± SEM in each group. Data were analysed by one-way ANOVA. Statistical differences associated with the Infected group were marked according to the colour of the symbol of each group. ANOVA, analysis of variance; Art, artesunate; ECM, experimental cerebral malaria; RMCBS, rapid murine coma and behaviour scale; SEM, standard error of mean; TMP, tetramethylpyrazine.

ECM mice (15.68 g) and TMP-treated mice (14.82 g) showed significantly lower weight on d7 p.i. as compared with the weight of mice in the Art (18.72 g) or Art + TMP treatment group (18.10 g), even until on d12 p.i. (Fig. 2B). The weight of mice in the Art + TMP group (18.05 g) was found to be higher than that of mice in the Art group (17.63 g) from d8 p.i. to d12 p.i. (Fig. 2B).

There was no significant difference in the body temperature of mice between the TMP group and the Infected group. However, on d9 p.i. and d11 p.i., the body temperature of mice in the Art group (33.2°C, 33.13°C) and the Art + TMP group (35.4°C, 34.59°C) was significantly higher than that in the TMP group (30.65°C, 27.42°C) and Infected group (29.48°C, 28.85°C) (*p* < 0.001). More noteworthy was the fact that the body temperature of mice in the Art + TMP group was significantly higher than that of mice in the Art group on d9 p.i. and d11 p.i. (*p* < 0.01, *p* < 0.05) (Fig. 2C).

### Art-TMP combination enhances the exploratory locomotion in ECM mice

The open field test was used to investigate the ability of autonomous activity (locomotion, exploration, and anxiety) in mice, as shown in Fig. 3A. The PbA-infected mice travelled shorter distances than control mice. Compared with the ECM mice (395.63 cm), the mice in the Art group and the Art + TMP group could travel longer distances in the open field experiment (845.82 cm for Art, 1081.59 cm for Art + TMP), which is consistent with our assessment of mice status using RMCBS. Notably, the results showed significant differences between mice in the Art and Art + TMP groups (Fig. 3A). Similarly, the total movement time that mice in the Art (75.02 s) and Art + TMP groups (82.13 s) spent was significantly longer than the time spent by mice in the Infected group (36.12 s) (Fig. 3B). Although significant differences in the distance travelled and the time spent in the central area were observed between mice in the Art, Art + TMP and Infected groups (Fig. 3C, 3D), all mice in the four groups did not differ in the percentage of distance travelled in the centre of the open field (Fig. 3E), indicating that mice that travelled longer distances and spent more time in the central area did not show an anxiety-like state.

**Figure 3.**
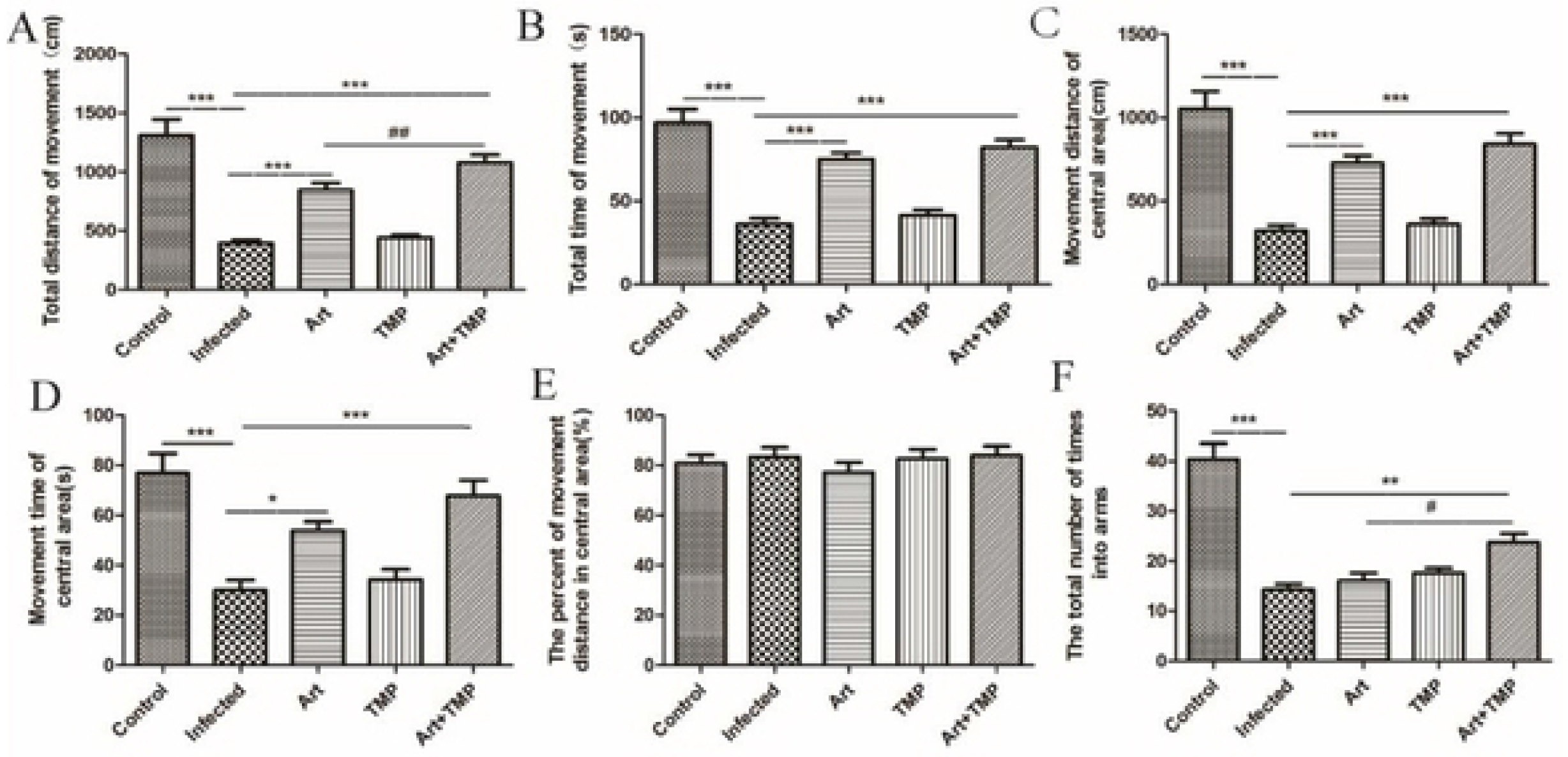
Improved autonomous activity in Art + TMP mice as measured by the open field test and Y-maze spontaneous alternation test. All PbA-infected mice with ECM were treated with drugs as described above. (A) Total distance of movement in the open field test on day 7 p.i. (B) Total time of movement in the open field test on day 7 p.i. (C) Distance travelled in the central area. (D) Time spent in movement in the central area. (E) The percentage of movement distance in the central area. (F) Effect of Art, TMP, Art-TMP combination drugs on number of entries in the closed arms following 8 minutes of maze exploration on day 8 p.i. in the Y-maze test. Data are expressed as mean ± SEM of two independent experiments. (n = 10 in control group and n = 13 in other groups). Art, artesunate; ECM, experimental cerebral malaria; PbA, Plasmodium berghei ANKA; SEM, standard error of mean; TMP, tetramethylpyrazine.

To study the effectiveness of new treatments on CM in animal models, animals were tested for spontaneous alternation in the Y maze. All drug-treated mice performed to similar extent in this test. The total number of times into arms (±SEM) was 16.00 times for Art, 17.62 times for TMP and 23.75 times for the Art + TMP mice. Consistent with stronger autonomous activity, Art + TMP mice spent significantly more time into the arms of the maze (23.75 times vs. 16.00 times) compared with the Art mice (Fig. 3E). It is suggested that the Art + TMP combination could increase the spontaneous alternation of mice and enhance the ability of autonomous activity and exploration to the novel environment.

### Art-TMP combination reduces cerebrovascular pathology and increases vessel integrity and patency

We next checked the hypothesis that the Art-TMP combination had a more profound effect than the Art and TMP alone on brain pathology. We found significantly fewer adherent leukocytes in mice in the Art + TMP group than in the untreated mice on d7 p.i. (Fig. 4A). In addition, MRI is considered as a valuable visualisation tool for tracking microvascular pathological changes *in vivo* [24]. To investigate whether the Art-TMP combination could improve vascular damage, mice in the four groups (*n* = 6) were measured by MRI. More abundant and unobstructed blood vessels were observed in mice in the Art + TMP group as compared with the dramatic reduction of vessel integrity and patency in untreated PbA-infected mice (Fig. 4B), which is consistent with our pathological results using HE staining. We next evaluated the changes in the expression levels of adhesion molecules in brain blood vessels, and found that the expression of ICAM-1 and VCAM-1 was significantly decreased in the Art + TMP group mice as compared with expression of these molecules in the ECM mice (Fig. 4D, E, *p* < 0.001). There was still a significant difference in the number of ICAM-1- and VCAM-1-positive vessels between mice in the Art + TMP and Art alone groups (Fig. 4D, 4E, *p* < 0.05, *p* < 0.01). In summary, these results showed that the overall protective effect of Art + TMP treatment was to help relax and widen the blood vessels and alleviate pathological damage.

**Figure 4.**
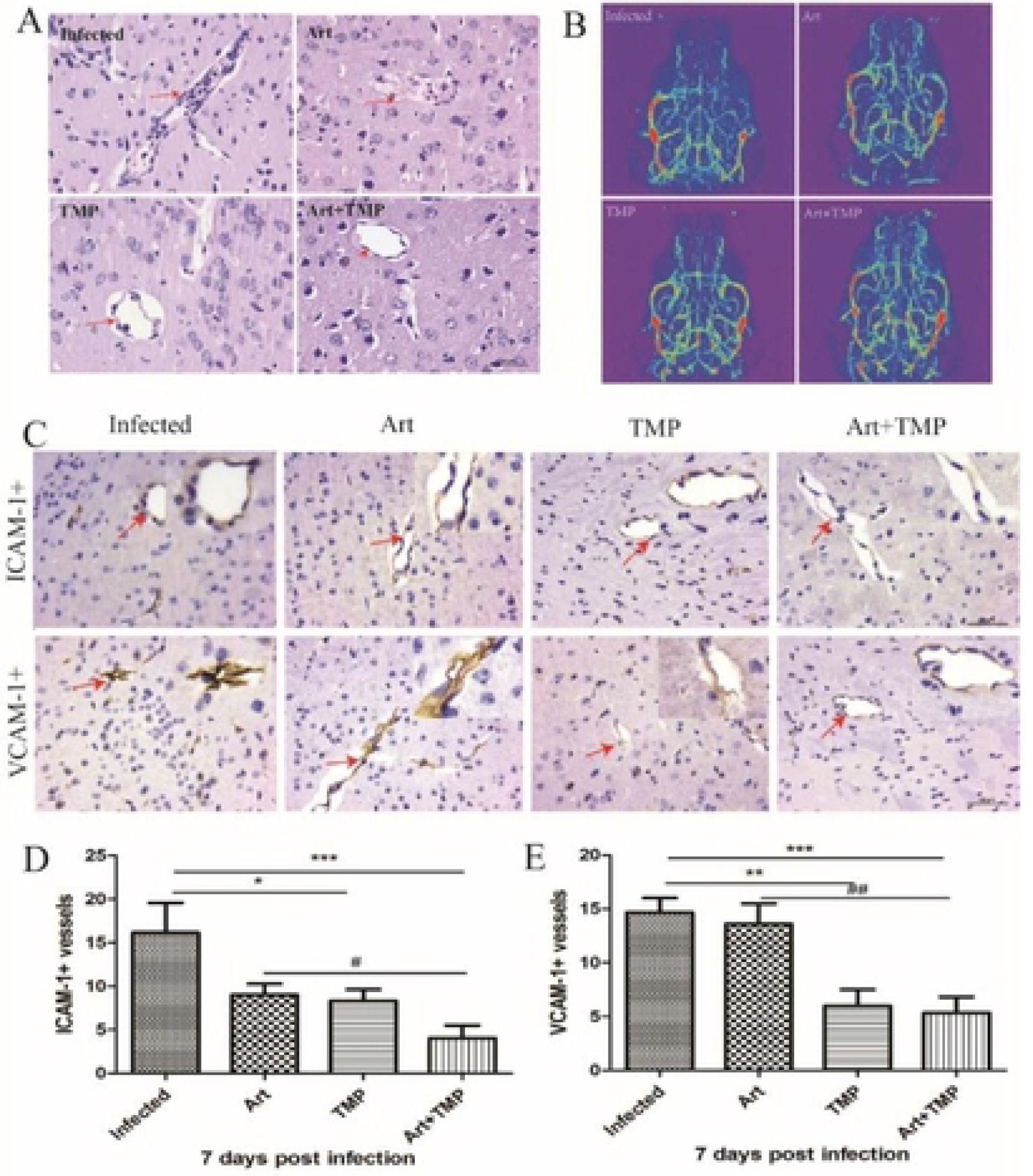
The efficacy of the Art-TMP combination was checked in reducing the cerebrovascular pathology and maintaining vascular integrity and patency in ECM mice. Inhibition of inflammation in the brain of PbA-infected mice after drug treatment as described above. (A) HE staining of the brain tissue sections. Arrows show the adhesion of leukocytes and pRBC to the blood vessels. All sections were stained with H&E (magnification, ×400). (B) Representative MRI images of the brain obtained by TOF-2D-FLASH scanning. (C) Representative images of ICAM-1- and VCAM-1-positive blood vessels (ICAM-1+, VCAM-1+) obtained by immunohistochemistry of the brain slices, with the arrow pointing in the direction of the microvessel. Upper right is the enlarged image of the vessel region. (D) Random ICAM-1+ blood vessels and (E) VCAM-1+ blood vessels (n = 20). Each mouse was randomly selected from 20 fields to evaluate the number of vascular expressions of ICAM-1+ and VCAM-1+ from 6 mice per group. Scale bar: 50 μm. Data are presented as mean of each group ± SEM. Data were analysed by one-way ANOVA. ANOVA, analysis of variance; Art, artesunate; ECM, experimental cerebral malaria; HE, haematoxylin and eosin; ICAM, intercellular cell adhesion molecule; MRI, magnetic resonance imaging; PbA, Plasmodium berghei ANKA; SEM, standard error of mean; TMP, tetramethylpyrazine; TOF, time-of-flight; VCAM, vascular cell adhesion molecule.

### Art-TMP combination inhibits activation of astrocytes in the cortex and maintains neuronal vitality in hippocampus

Activation of astrocytes and microglia acts as a crucial player in several pathophysiological changes in the central nervous system (CNS) [25]. Glial fibrillary acidic protein (GFAP) is a key protein expressed in astrocytes and is significantly up-regulated when nerves are damaged [26].We assessed astrocyte activation using the known astrocyte marker GFAP. The number of GFAP-positive cells in the brains of mice in the Art + TMP group significantly decreased as compared with that in mice in the Infected group and Art group (Fig. 5B, *p* < 0.05, *p* < 0.01). Immunohistochemistry using anti-NeuN antibody was performed to assess neuronal viability [27]; the number of NeuN-positive cells significantly increased in the brains of mice in the Art + TMP group as compared with that in mice in the Art, TMP-treated and Infected mice.

**Figure 5.**
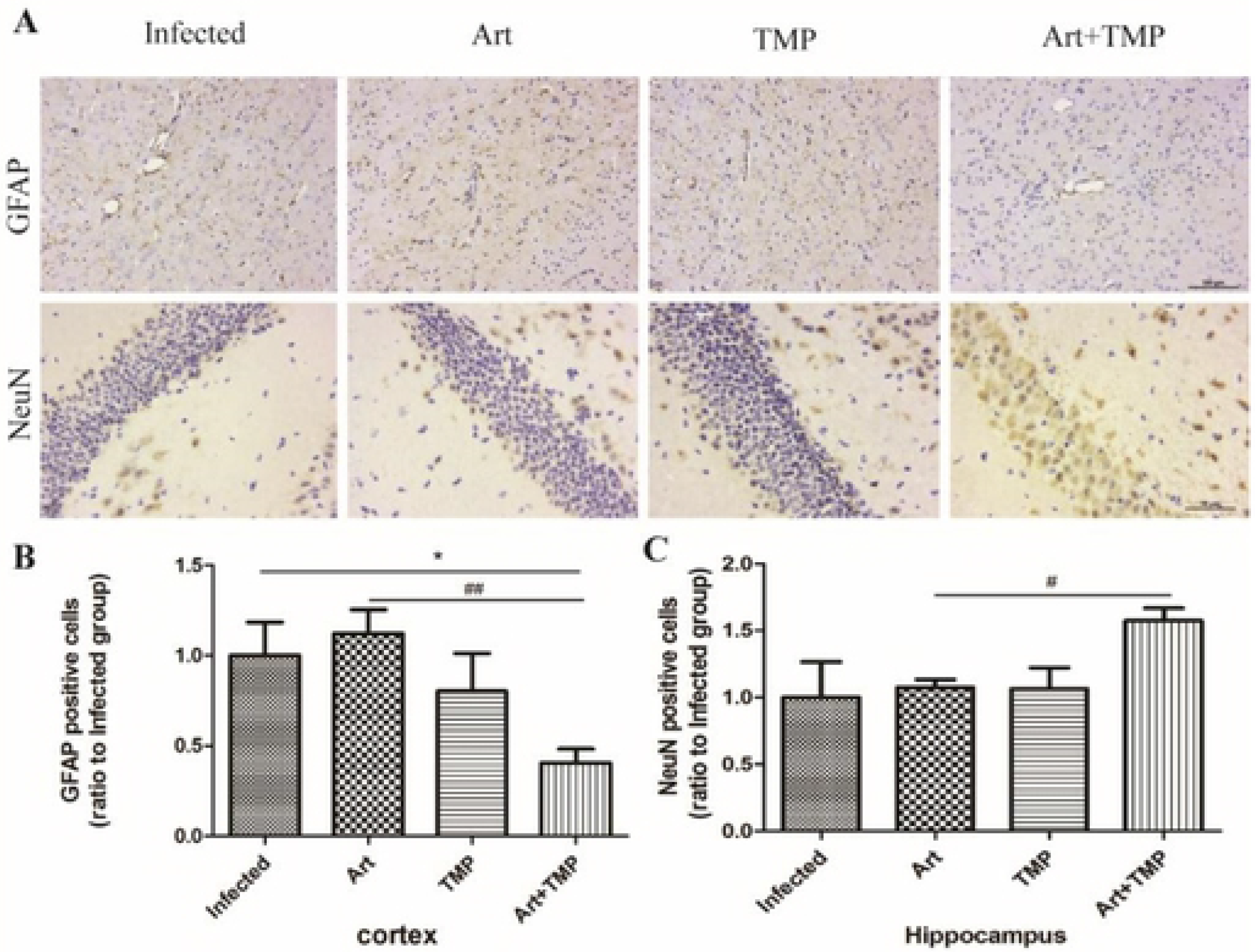
The Art-TMP combination suppresses astrocytes activation and sustains neuronal viability. Immunohistochemistry using anti-GFAP and anti-NeuN antibodies was performed in the brain of ECM mice after the treatments as described above. (A) Representative images of immunostaining of GFAP in the cortex and of NeuN in the hippocampus. (B, C) Quantitative analyses of GFAP-positive cells in the cortex and NeuN-positive cells in the hippocampus. Results were calculated as ratio to the Infected group and are expressed as means ± SEM (n = 6 mice in each group). Scale bar: 100 μm. ECM, experimental cerebral malaria; GFAP, glial fibrillary acidic protein; NeuN, neuron-specific nuclear protein; SEM, standard error of mean; TMP, tetramethylpyrazine.

### Modulation of cytokine production by Art-TMP combination in ECM mice

Cognitive deficits have been reported to be associated with changes in neurotrophic factors [28], including BDNF and nerve growth factor (NGF). Considering that neurological dysfunction of CM is associated with cognitive impairment, we studied the changes in the concentration of neurokines in mice treated with drugs. Our results showed that mice in the Art + TMP group showed significantly elevated levels of BDNF and NT-3 as compared with mice in the infected group (Fig. 6A, 6B, *p* < 0.01). The secretion of growth factors can promote endothelial cell proliferation and neovascularisation in the perivascular region, resulting in tissue repair. Our study revealed that the levels of b-NGF and VEGF-A in mice in the Art + TMP group were higher than those in mice in the Infected group. Interestingly, these were significantly higher than those in the Art group (Fig. 6C, 6D, *p* < 0.001). In addition, levels of pro-inflammatory factor TNF-α were also significantly lower in mice in the Art and Art + TMP groups than those in mice in the Infected group (Fig. 6E, *p* < 0.01).

**Figure 6.**
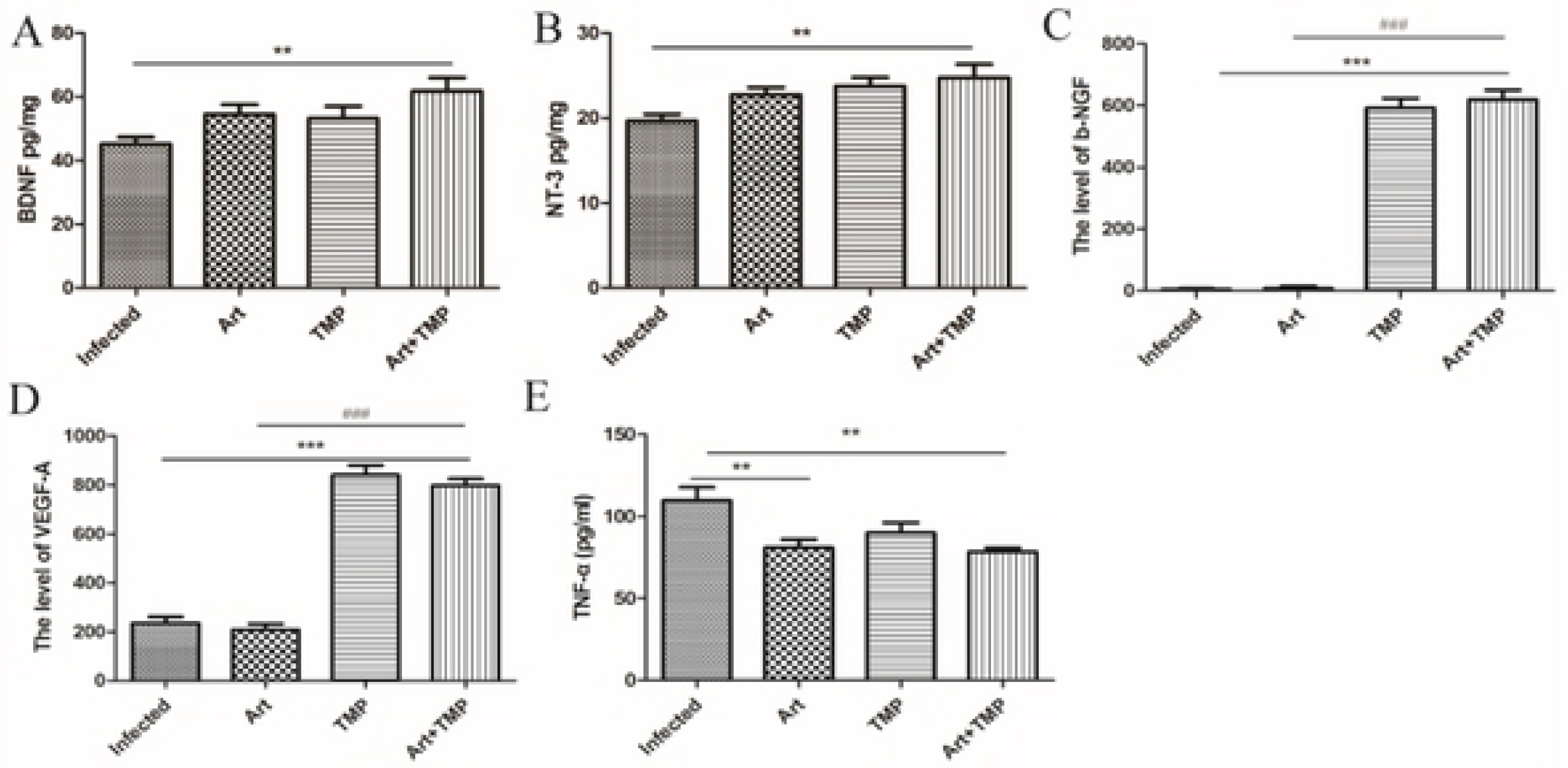
Expression levels of BDNF, NT-3, b-NGF, VEGF-A and TNF-α in the brain tissue of ECM mice after treatments as described above (n = 6 in each group). Data are expressed as mean of each group ± SEM. Data were analysed by one-way ANOVA. ANOVA, analysis of variance; BDNF, brain-derived neurotrophic factor; ECM, experimental cerebral malaria; NT-3, neurotrophic factor-3; NGF, nerve growth factor; SEM, standard error of mean; TMP, tetramethylpyrazine; TNF, tumour necrosis factor; VEGF, vascular endothelial growth factor.

### Proteomic analysis: differentially regulated proteins among treatments

Both Art and TMP are known to improve the neurological symptoms of ECM mice. We therefore performed a proteome analysis to obtain further information about the protein profiles in the presence of Art alone, TMP alone and Art-TMP combination. Among the four treatments, a total of 5,324 proteins were identified. We screened 192 differentially expressed proteins (DEPs) following treatment with Art; of these, 128 and 64 proteins were found to be down- and up-regulated, respectively. There were 41 up-regulated DEPs and 48 down-regulated DEPs in the TMP group and 177 up-regulated and 217 down-regulated proteins in the Art + TMP group versus the ECM group. A total of 142 and 99 proteins were found to be up- and down-regulated in the Art + TMP group and the Art group, respectively. The Art + TMP versus the TMP group included 133 up-regulated and 107 down-regulated DEPs. These DEPs comprised 12 and 16 up- and down-regulated proteins, respectively, overlapping among treatments. Detailed information can be found in the S1 Table and S2 Table. The top 10 most significant brain DEPs in the Art, TMP and Art + TMP groups can be found in the S3 Table. We then compared the brain proteins that increased in mice in Art, TMP, and Art + TMP groups relative to those in the Infected mice. As illustrated by the Venn diagram in Fig. 7A, a set of 12 proteins (ANR63, NRP2, RGS9, PENK, PLD2, BMP1, LIGO2, EPHA5, CC85B, NCKX4, CHAP1, and NASP) were elevated in all drug groups. As shown in Fig. 7B, levels of a set of 16 proteins (MYL1, NFL, PERI, NFH, LPAR1, H32, K22E, MYH4, K1C15, ARMD3, PCP, K2C79, RFA3, MTNA, PP4P2, and S22A4) were found to be reduced in all drug groups. Proteins unique to the Art + TMP group included those belonging to the neuroprotective pathway (SEMCB1, HAX1, CD5R1, HPCA, GARE1, CD166, PC4L1, TIAM2, ROBO2, and SLIT2) and cerebrovascular protection pathway (BAIP3 and PCP4L1).

**Figure 7.**
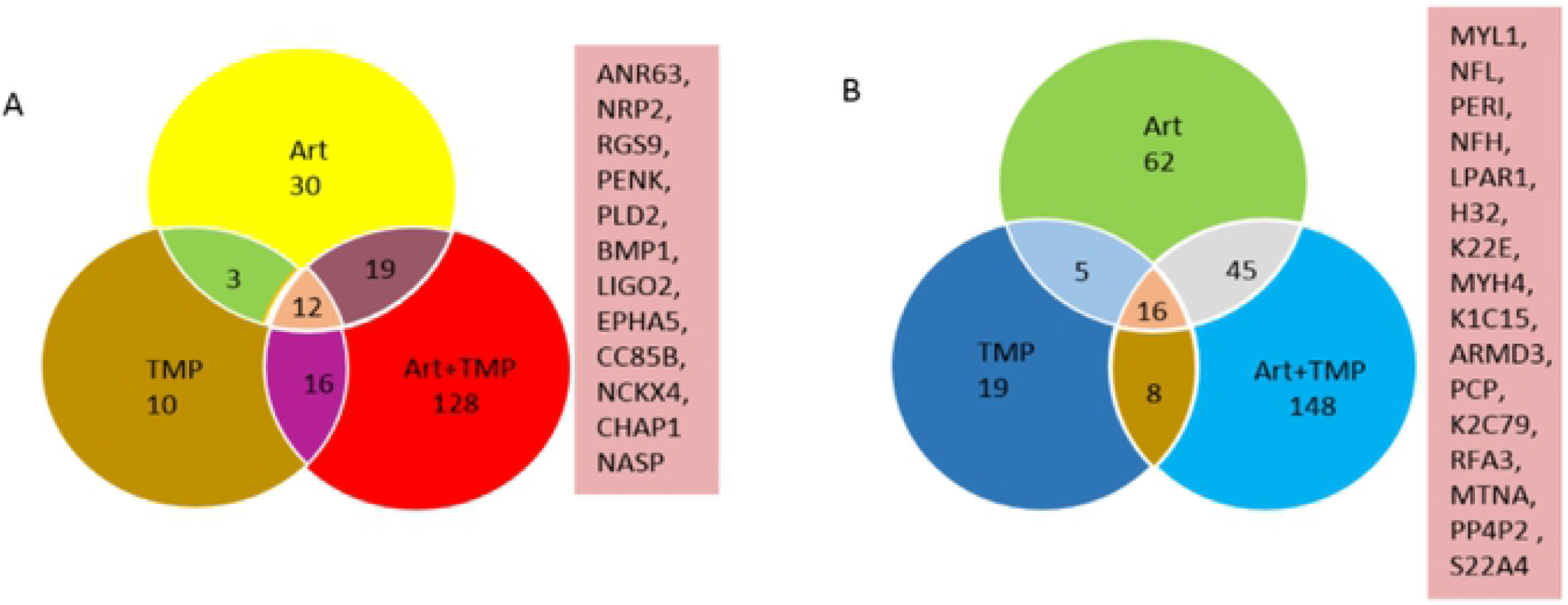
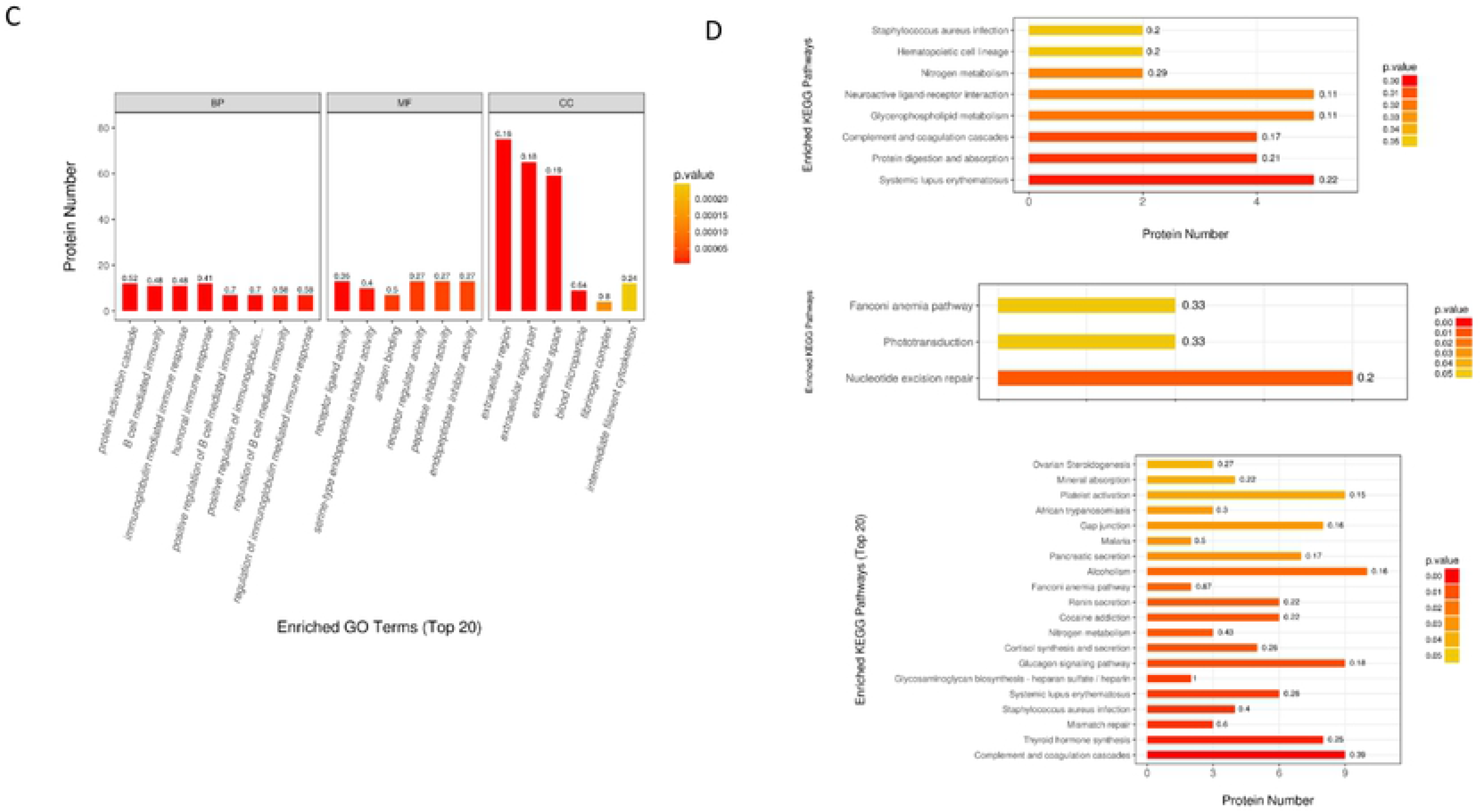
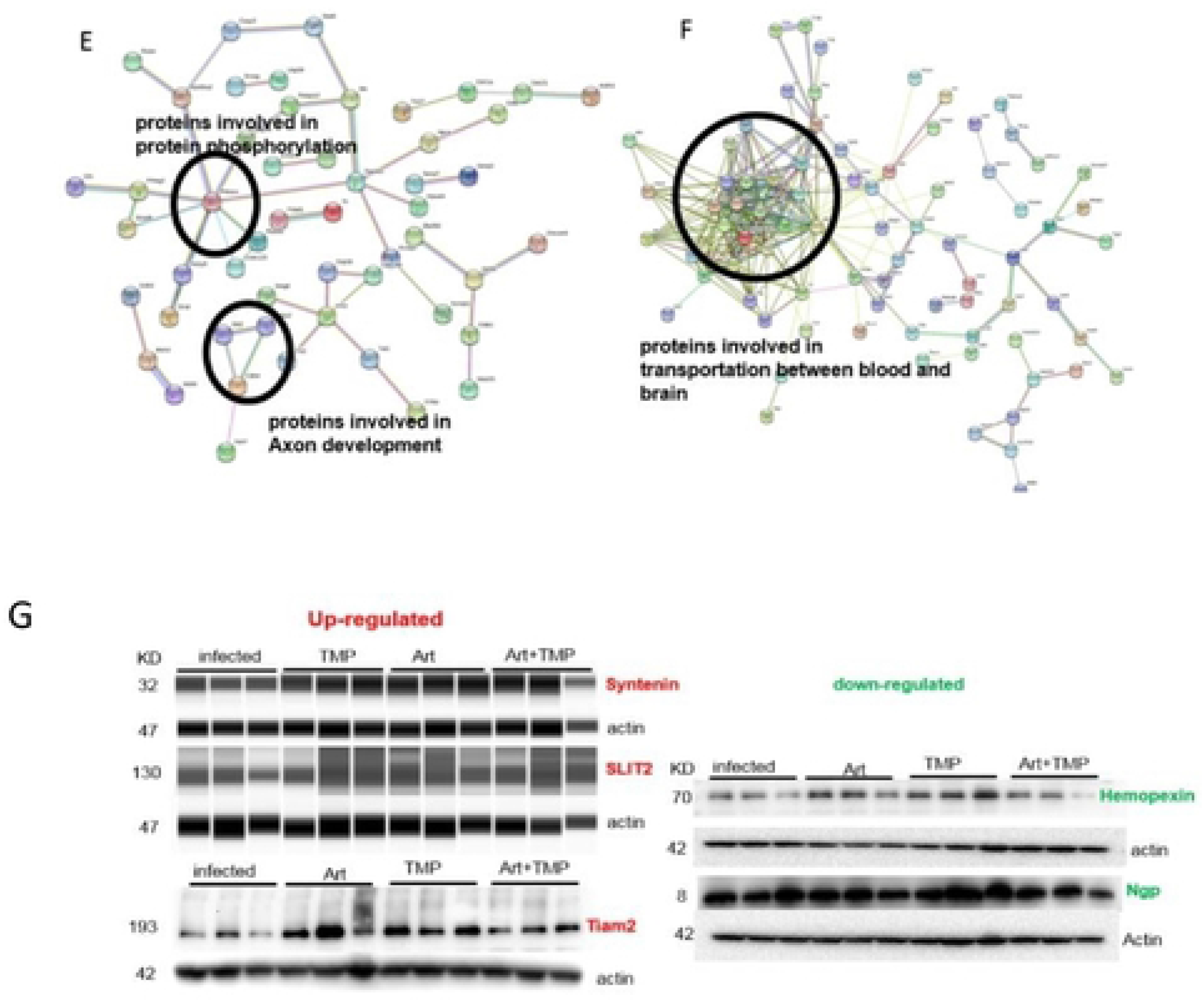
iTRAQ proteomic analysis of brain tissues in ECM mice after different drugs treatment. (A) unique up-regulated proteins overlapping among Art, TMP and Art-TMP combination treatment relative to cerebral malaria mouse. Protein names for overlapping up-regulated proteins are shown to the right of the Venn diagram. (B) unique down-regulated proteins overlapping among Art, TMP and Art-TMP combination treatment relative to cerebral malaria mouse. Protein names for overlapping up-regulated proteins are shown to the right of the Venn diagram. (C) GO annotation of differentially regulated protein (ratio: > 1.2 or < 0.8) functions. Y-axis represented the number of identified proteins in each GO category. (D) KEGG pathway analysis. KEGG pathway enrichment analysis results for comparison Art (differentially expressed proteins in Art treatment group compared with ECM mice group), for comparison TMP (differentially expressed proteins in TMP treatment group compared with ECM mice group) and comparison Art+TMP (differentially expressed proteins in Art+TMP treatment group compared with ECM mice group) are shown. The y-axes indicates the significantly enriched KEGG pathway;The x-axes represent the number of differentially expressed proteins contained in each KEGG pathway (the P value is calculated based on Fisher’s exact test). The color gradient represents the magnitude of the P value and the tab above the bar shows the enrichment factor, indicating the number of differentially expressed proteins involved in a KEGG pathway as a percentage of the number of proteins involved in the pathway in all identified proteins. (E and F) The protein-protein interaction (PPI) network of proteins in Art+TMP treated groups based on STRING analysis. A total of 127 differentially up-regulated proteins and 148 differentially down-regulated proteins are shown in E and F PPI network, respectively. Strong associations are represented by thicker lines. (G) Validation of the proteomics results. Western blotting analysis of five proteins selected from the proteomics data. Art, artesunate; ECM, experimental cerebral malaria; TMP, tetramethylpyrazine.

**Figure 8.**
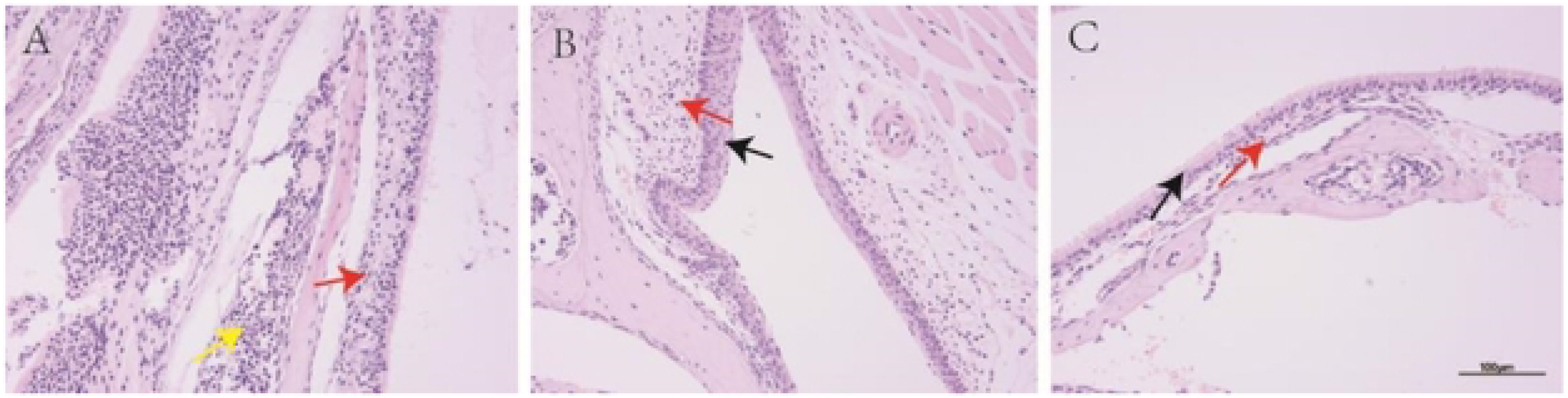
Histopathological sections of HE staining of the nasal mucosa of healthy C57BL/6J mice treated with Art + TMP combination (20 mg·kg-1) by intranasal administration. (A) A control group received intranasal administration of placebo solution. (B) C57BL/6J mice received intranasal administration of Art + TMP once. (C) C57BL/6J mice received intranasal administration of Art + TMP four times. Scale bar: 100 μm. Red arrows indicate inflammatory cell infiltration; leukocyte aggregation can be seen in the lumen, as shown by the yellow arrow. The epithelium of the nasal mucosa is intact and cilia on the epithelium are clearly visible, as shown by the black arrow. Art, artesunate; ECM, experimental cerebral malaria; HE, haematoxylin and eosin; TMP, tetramethylpyrazine.

### Functional enrichment analysis

We observed more up- and down-regulated proteins in the Art-TMP combination treated mice than in mice treated with either Art or TMP individually, which is consistent with improved effects exerted by Art-TMP combination administration than Art or TMP treatment alone in ECM mice. Therefore, we focused on DEPs specific to the Art-TMP combination group to perform GO and KEGG enrichment analyses to obtain more information about the mode of action of the combination therapy. GO term enrichment revealed that most DEPs were involved in protein activation cascades, B cell-mediated immunity and its regulation, immunoglobulin-mediated immune response and its regulation, humoral immune response and positive regulation of immunoglobulins. Detailed information is provided in Fig. 7C. Notably, functions of DEPs in the Art, TMP, and Art-TMP combination groups mainly involved binding, catalytic activity, regulation of molecular functions, transporter activity, and structural molecules activity; these are majorly involved in cellular processes, biological regulation, metabolic processes, and response to stimuli. We annotated all identified DEPs of Art, TMP, and Art-TMP combination groups using the KEGG database and mapped these to 166, 72, and 234 KEGG pathways, respectively. 8, 3 and 20 pathways in Art, TMP, and Art-TMP combination treatment groups were considered statistically significant, respectively, including those involved in systemic lupus erythematosus, protein digestion and absorption, complement and coagulation cascades, glycerophospholipid metabolism, neuroactive ligand–receptor interaction, nitrogen metabolism, haematopoietic cell lineage, and *Staphylococcus aureus* infection in the Art group (Fig. 7D-Art); nucleotide excision repair, phototransduction, and Fanconi anaemia pathway in the TMP group (Fig. 7D-TMP); complement and coagulation cascades, thyroid hormone synthesis, mismatch repair, *S. aureus* infection, systemic lupus erythematosus, glycosaminoglycan biosynthesis-heparan sulfate/heparin, glucagon signalling pathway, cortisol synthesis and secretion, nitrogen metabolism, cocaine addiction, renin secretion, Fanconi anaemia pathway, alcoholism, pancreatic secretion, malaria, gap junction, African trypanosomiasis, platelet activation, mineral absorption and ovarian steroidogenesis in the Art-TMP combination group (Fig. 7D-Art+TMP). Thus, there are more KEGG pathways involved in the Art-TMP combination treatment group than in the Art or TMP treatment groups, similar to the results obtained for DEPs and GO enrichment analysis in the three drug groups. We speculate that the Art-TMP combination treatment may improve the neurological symptoms in CM mice by interfering with the neuroactive ligand–receptor interactions, transporters, and certain metabolic pathways. Of particular interest are the DEPs, GO, and KEGG analyses that revealed more significant changes in the Art-TMP combination treatment group than in the Art and TMP treatment groups. With these results, we next focused on analysing DEPs only in the Art-TMP combination treatment groups, attempting to find the mechanisms that could explain the improvement of symptoms observed in ECM mice after Art-TMP combination treatment.

To better explore the functional relationships among DEPs, a network was constituted using protein–protein interactions of the significantly up-regulated and down-regulated proteins in the Art-TMP combination group, as shown in Fig. 7E and 7F, respectively. A group of significantly up-regulated proteins was found to actively interact; this included proteins involved in protein phosphorylation: Prkaca, Rab8a, Adcy5, prkag2, Rps6ka2, Rap2b, and Ppp3cc, and proteins involved in the axon development: Slit2, Robo2, and Cdh4. A group of significantly down-regulated proteins was also found to actively interact and included proteins involved in the transportation between blood and brain: Fgg, Apoh, ltih4, Hrg, Fga, Fgb, Orm1, Fn1, Serpina3k, alb, Apoa1, Hpx, serpina1b, Mug1, and Fetub. Based on these findings, we predicted that these different proteins may be involved in improving neurological symptoms of CM in mice, thereby acting as novel candidates for Art-TMP combination treated ECM mice.

### WB validation of iTRAQ analysis

To verify the accuracy of the data, the selected differential proteins were validated at the protein level by western blotting (WB) (Fig. 7G). Considering the neuroprotective effects of Art and TMP on CM, we studied the levels of proteins that demonstrated significant cardiovascular protection and neuroprotection of the brain tissue proteome in the presence of drugs.

We selected five proteins including three up-regulated and two down-regulated proteins in the Art+TMP group to validate Syntenin, Slit2, Tiam2, Emopexin and Npg proteins, based on their statistical significance and biological relevance with respect to improvement seen in CM. When measured by WB or WES, the significance of Syntenin, Slit2 and Tiam2 proteins was validated on an independent set of brain samples. Increased levels of Syntenin, Slit2 and Tiam2 proteins in Art, TMP and Art+TMP groups were significantly higher than those in the Infected group. The levels of hemopexin and Npg proteins were significantly lower in Art, TMP and Art+TMP groups as compared to their levels in the Infected group, which were consistent with the iTRAQ analysis.

### Intranasal administration of artesunate combination does not affect olfactory bulb tissue

Intranasal administration of healthy C57BL/6J mice with Art-TMP combination (20 mg·kg^−1^) or placebo solution did not result in local irritation symptoms (such as asthma, cough, vomiting, asphyxia, etc.). Also no abnormalities (such as breathing, exercise and behaviour) were observed in animals. The safety of administration was evaluated by HE-stained histopathological sections of the olfactory bulb tissue. Compared with placebo-treated mice, the treated mice did not show severe nasal lesions, with only partial vasoconstriction in the connective tissue of the mucosa and slight lymphocyte accumulation in the lumen. However, this pathological change also existed in placebo-treated mice, indicating that it was unrelated to intranasal administration of Art-TMP combination.

## Discussion

Cerebral malaria is the most virulent and deadliest clinical manifestation of malaria. In the Sub-Saharan Africa, five-year-old or younger children are the most susceptible to developing CM; the infection is associated with 90% mortality in this region [29]. Tragically, one in four children who recovers from CM suffers from neurological deficits, including hemiplegia, cerebellar ataxia, hypotonia, paralysis, aphasia, behavioural disorders and attention deficit [6, 30-33], indicating that elimination of the parasite does not completely improves the clinical outcomes of the infection. Thus, developing a safe, effective, and novel adjunctive therapy is the need of the hour to counter the disease.

Pathophysiology of CM is still controversial. Sequestration of pRBCs and immune cells into the brain vasculature leads to vascular obstruction, hypoperfusion and hypo-oxygenation that are considered to be the major contributors to CM. Clinical evidence from patients with CM suggests fibrinogen levels to be elevated in the brain parenchyma near cerebral blood vessels that are filled with pRBCs along with axonal injury associated with haemorrhage and demyelination [29, 34, 35]. Another clinical study on CM treatment in Cambodia reported that RBCs infected with trophozoites accumulated in the blood vessels of patients resulting in ischaemia and hypoxia in the brain, consequently causing irreversible neurological damage. Even if the patients were administered artemisinin and heparin, tiny blood vessel obstructions remained[36]. Therefore, it is essential to explore a method of timely clearance of the microvascular occlusion to alleviate and eliminate cerebral ischaemia and hypoxia injury in patients with CM and improve long-term neurological deficits and dysfunction. In the present study, we used an ECM mouse model to investigate the potential of TMP as an adjunctive therapy to improve the neurological symptoms and survival. Results of our study revealed lower parasitaemia, better clinical outcomes, improvement of histological and immunological features in ECM mice after treatment with TMP + Art as compared with ECM mice. Subsequently, we performed quantitative proteomic analysis to compare the brain proteome profiles of ECM mice treated with Art, TMP and Art + TMP to explore the possible targets of Art + TMP in treating ECM.

Firstly, we successfully replicated the ECM model that exhibited neurological damage including hemi- and paraplegia, ataxia, and convulsions, consistent with previous studies [37, 38]. ECM mice were treated by intranasal administration with Art and TMP at d2 to d5 p.i. to investigate the effect of Art-TMP combination in preventing the incidence of ECM at an early stage of malaria infection. Several researchers have initiated empiric novel treatments of ECM administered before, on, or just after the first day of infection for evaluating various effects at different stages of malaria infection [39–43]. Our future studies aim to study the drug effect after 3 to 7 days of infection on long-term prevention of neurological damage in ECM mice. In the present study, we did not observe any significant changes in parasitaemia, body weight and survival status of ECM mice after TMP treatment alone. However, we observed a considerable delay in death from ECM after TMP treatment from d8 p.i. to d9 p.i. Moreover, mice in the Art and Art-TMP combination groups exhibited prolonged survival (even close to 100%) in the Art-TMP combination group, as well as a decrease in parasitaemia, and an increase in body weight and temperature. More importantly, a significant decrease in the proportion of large trophozoite stage of *Plasmodium* in the blood smear of the two groups was observed. Notably, a lower parasitaemia and increased body weight and temperature were observed upon treatment of infected mice with Art-TMP combination than those treated with Art alone, suggesting that the Art-TMP combination treatment could successfully reduce the pathological outcomes and improve the clinical signs in ECM mice.

One of our previous study has confirmed that the Art-TMP combination treatment significantly improved the cerebral vascular occlusion in ECM mice than in mice in the Art-treated group [44]. Same results were also observed in the present study, especially the significant improvement of symptoms including decreased adhesion of parasite-infected erythrocytes and immune cells to cerebral microvessels after TMP monotherapy, which had the same effect as the Art-TMP combination treatment. Indeed, abnormal microvascular integrity and cerebral perfusion were observed by MRI in ECM mice, consistent with the findings of occlusion effect of brain microvasculature in ECM mice from HE studies. We observed that Art and TMP monotherapy could improve the above-mentioned symptoms, with the Art-TMP combination therapy showing the most significant improvement by MRI. Activation of endothelial cells is manifested in several ways, for example increased expression of adhesion molecules. ICAM-1 (CD54) and CD36 are two major binding partners for the PfEMP1 protein on PfRBCs that contribute to PfRBC sequestration. The important factor of ICAM-1 expression in the development of CM is well established. Tripathi *et al.* found that exposure of the human brain microvascular ECs to PfRBCs induced the expression of ICAM-1 [45]. Favre *et al.* reported that ICAM-1-deficient mice were protected from CM [46]. Interestingly, our study found that TMP monotherapy and Art-TMP combination therapy significantly reduced the activation of brain microvascular endothelial cells with a lower expression of ICAM-1 and VCAM-1 in the endothelial cells of ECM mice than that in the untreated ECM model mice. However, no significant effect of Art monotherapy on endothelial activation among ECM mice was observed. These results demonstrate that Art-TMP combination plays a major role in defining the pathological outcomes and clinical signs in ECM mice. Specifically, Art exerts a direct killing effect on *Plasmodium* parasite. TMP may play roles in decreasing endothelial activation, reducing sequestration of iRBC and adhesion of leukocytes, and increasing blood perfusion in the brain, whereas a combination of the two drugs exerts a potent synergistic effect on mouse survival, parasitemia, and body weight and temperature.

RMCBS can evaluate the related behavior of mice to reflect the real-time function of central nervous system, it can be used to objectively evaluate the disease process of mice and provide a tool for evaluating new adjuvant therapies. During the observation period, ECM mice gradually showed signs of walking instability, ataxia, fur curl, arch back, decreased toe response and disappearance of auricle reflex, convulsions, coma symptoms and death. The results of RMCBS score were consistent with those of mice. In this study, significant improvement in the RMCBS scores of ECM mice was observed in Art-treated group. Moreover, the Art-TMP combination also significantly improved the RMCBS scores of mice at d9, 10, 11 p.i. as compared with the Art group, suggesting that Art-TMP combination exerted synergistic effects on improving coordination, exploration, muscle strength, reflex, self-protection and health behaviour of ECM mice. Based on the ECM-specific neurological and behavioural evaluation, we confirmed that the combination of Art and TMP prevented significant deterioration of neurological functions. Furthermore, we evaluated whether this combination could improve cognitive and behavioural changes in ECM mice, and further investigated the effects on long-term neurological dysfunction in CM.

The open field test can qualitatively and quantitatively monitor the spontaneous activity of experimental animals. Our results showed that the Art alone and the Art-TMP combination changed the total moving distance and total movement time of ECM mice, which is in agreement with the results of RMCBS score, suggesting that the Art therapy and Art-TMP combination have a definite impact on the athletic ability and exploring behaviour of ECM mice. The results of the Y maze test also indicated that the Art group and Art + TMP combined group significantly affected the total number of arms and improved the behaviour function to a certain extent, suggesting that the adjunctive therapy could protect against ECM-induced behaviour impairment.

Notably, previous studies have implicated anxiety-like behaviour to occur in ECM mice in the open field experiment[31]. However, in our ECM model, we did not observe any increase in the activity of mice in the central area of the empty field and other anxious activities. We did observe a significant difference in the activity of the central area of the empty field. A possible explanation for this could be that the ECM mice model established in our laboratory mainly exhibited degenerative changes in spontaneous activity and exploring behaviour, thus showing contracture, apathetic, reduced activity, reduced response to the stimulus, and gradually appeared chest, hemiplegia and convulsions symptoms. The overall state of our mice was different from the anxiety-like changes reported in the literature; however, a combination of the Art + TMP revered these effects in different regions of the empty field. The TMP (alone) administration had no significant impact on the relevant indicators of exercise, exploration, and cognition.

We next confirmed the synergistic effect of the Art-TMP combination treatment on neurobehavioural signs in ECM mice with a further exploration of the neuroprotective efficacy in ECM mice. Astrocytes are common CNS-residing cells that are essential for regulating the blood flow and maintaining the blood–brain barrier, thus maintaining the immune defences in the CNS. Alteration of the cerebral microcirculation is an important factor in the pathogenesis of CM. Sequestered PfRBCs interact closely with the cerebrovasculature, enhancing permeability, endothelial activation, and vascular obstruction, thereby contributing to cerebral microcirculatory alterations. As shown by our HE staining results, large trophozoite parasite pRBCs and WBCs adhered to the brain microvessels of ECM mice, leading to ischaemia and hypoxic injury. Trophozoite-mediated occlusion of microvessels also activated the glial cells (Figs. 4 and 5). After drug administration, the number of GFAP-positive cells in the cerebral cortex of mice in the Art-TMP combination group significantly reduced. Moreover, the abnormal activation of astrocytes was decreased. The Art-TMP combination administration also enhanced the expression of neuronal NeuN in the hippocampus, suggesting the neuroprotective effects of this combination against neuronal damage in ECM mice. TMP and Art-TMP combination groups could significantly increase the expression levels of BDNF, NT-3 and bNGF in the brain tissue, indicating the neuroprotective effect to be related to the increasing supply of neurotrophins and enhanced neuronal repair and regeneration. Both single TMP administration and Art-TMP combination elevated the expression of VEGF-A, suggesting that TMP treatment could promote the proliferation of endothelial cells and the formation of new blood vessels surrounding the site of injury, thus restoring the blood supply to the ischaemic tissue and diminishing the harmful effects of ischaemia on the brain tissue. Interestingly, single Art treatment had no significant effect on the above signs, suggesting its inability to relieve the complications of nerve damage caused by microvascular occlusion in ECM mice even after killing of the *Plasmodium* parasite. However, the TMP treatment of ECM mice improved the above conditions via improving the nerve nutritional status, promoting neuronal and vascular tissue repair and regeneration, and other ways of restoring damaged neurological function in ECM mice. Given that Art alone was unable to improve the neurological protective function, the effect of single TMP on the complication of ECM above was consistent with the Art-TMP combination therapy with equal effectiveness. We speculate that the protective effect of Art-TMP combination therapy against neurological functional deterioration could be a major effect of TMP.

A synergistic intervention between Art and TMP reflects ‘division of labour’ in ECM mice. We used proteomics and bioinformatics to identify the potential synergistic mechanism of these two drugs. The present study identified the DEPs profiles before and after pharmacological treatments in ECM brain samples, followed by enrichment analysis and network analysis to discover the changes in proteins in response to drug administration. These analyses provided vital clues with respect to the mechanism of action of drug or biomarkers and toxicity that could guide future clinical trials. A total of 28 DEPs were identified to be significantly differentially expressed among the Art, TMP and Art-TMP combination groups. Our study found that there were more up- and down-regulated proteins in the Art-TMP combination treated mice than in mice treated with either of the individual drugs Art and TMP, a finding consistent with the therapeutic improved effects of the Art-TMP combination administration as compared to Art alone and TMP alone treatments in ECM mice. Therefore, we focused on studying DEPs specific to Art-TMP combination administration and performed GO and KEGG enrichment analyses. GO analysis revealed these DEPs to be associated with immune response and receptor regulator activity, whereas the KEGG analysis found these to be associated with platelet activation, African trypanosomiasis, malaria, nitrogen metabolism, systemic lupus erythematosus, and complement and coagulation cascades. The PPI network analysis of up- and down-regulated proteins of the Art-TMP combination group revealed differentially up-regulated proteins to be related to axon development. This included PRKACA protein that has been reported to be down-regulated and may be involved in the neuronal damage in patients with Parkinson’s disease [47]. However, our study found the PRKACA-centred protein pathway to be up-regulated in ECM mice after Art-TMP combination therapy, demonstrating this pathway to be involved in the molecular mechanism of action of drugs. Similarly, the PPI analysis revealed the down-regulated proteins specific to the Art-TMP combination group to be centred on the blood–brain transport, which plays important roles in the pathological improvement of ECM. Together, the bioinformatics studies predicted that the neuroprotective effects of the combined Art and TMP therapy may be associated with axon development or blood–brain transport.

We identified two down-regulated proteins, hemopexin and Ngp and three up-regulated proteins, namely Slit2, Tiam2 and Syntenin in the ECM mice brain tissues treated with drugs. Our WB validation analyses showed that the protein levels of Slit2, Tiam2, Syntenin, Hemopexin, and Ngp matched with the changes in the levels of proteins in the ECM mice by iTRAQ analysis after combined Art and TMP treatment. Up-regulation of hemopexin has been reported in the PbA-infected brain tissues as compared to the Pb NK65 (a *Plasmodium* strain that do cause ECM) and control mice [48]. Our research revealed reduced levels of hemopexin in ECM mice by the iTRAQ analysis, especially in the Art-TMP combination treated group by WB validation. These findings indicated that this protein may be involved in the ECM pathogenesis, and Art and TMP administration reversed pathological functions of hemopexin leading to a reversal of pathogenic neurological signs and enhancing the viability of neuronal cells in ECM mice.

We identified elevated levels of various complement components, immunoglobulin components, and factors involved in the coagulation cascade in ECM mice. Dysregulation of the coagulation system is a characteristic feature of severe malaria and impaired synthesis of clotting factors is a pathological reason behind it. Syntenin is one of the intracellular adaptor proteins that interacts with several proteins and regulates a number of pathways, such as immune-related pathways and those regulating angiogenesis and axonal growth [49, 50]. In our study, Syntenin was found to be elevated in ECM mice treated with Art-TMP combination, suggesting that complement and coagulation cascade pathways may be altered significantly as a consequence of up-regulation of the immune-associated proteins, such as Syntenin. This may also explain why neurological signs improved in the Art- and TMP-treated ECM mice. In addition to immune-related pathways, Slit–Robo signalling pathway also governs axon growth and angiogenesis. Interestingly, Slit2 protein was also up-regulated in the ECM mice brain after administration of Art and TMP, providing further evidence for the mechanism of action of these drugs that involves axon growth and angiogenesis.

Intranasal administration offers several advantages including high bioavailability, no liver first-pass effect, rapid absorption, and rapid onset of action of drugs that can easily enter the cerebrospinal fluid in the CNS, thus specifically targeting the brain. In ECM mice, a previous study reported the intranasal delivery of the anti-malarial drug Art to be an efficient way to contribute to decreasing malaria-related mortality [51]. Similarly, we treated ECM mice using Art or TMP or Art-TMP combination intranasally and observed the same results as reported by the above study; however, we also observed TMP to play a neuroprotective role in the ECM mice as a result of the targeted delivery to the brain.

## Conclusion

In summary, we have proposed an efficient combination treatment for CM employing Art and TMP, hopefully providing a potential effective adjuvant treatment for the clinic. Our study demonstrated the synergistic effect of Art-TMP combination treatment on ECM as compared to Art or TMP treatment alone. This protective effect is consistent with the roles of the two drugs in protecting the neuronal system and maintaining cerebrovascular integrity. The combination of Art and TMP increased the survival of ECM mice, prevented damage to the nervous system and improved clinical outcomes. These effects were associated with decreased cerebrovascular occlusion, increased expression of neurotrophic factors, increased blood flow to the damaged area, decreased expression of ICAM-1 and VCAM-1 in the brain endothelium, better integrity of the cerebrovasculature and reduced inflammatory factors in the ECM mice brain. These results indicated that Art-TMP combination could act as an effective therapy for CM with neuroprotective, anti-inflammatory and cerebrovascular integrity preservation effects. Despite the positive synergistic effects of the combination of Art and TMP, one caveat of our research is the use of a low dose of Art, which could only decreased the parasitaemia in the ECM mouse model. Further studies using curative doses of Art combined with TMP are warranted to mimic the situations in human CM. Considering the short-term therapy of TMP to be safe, clinical trials on TMP will help determine its potential as an adjunct treatment for human CM. To study the possible targets of Art-TMP combination in treating ECM, iTRAQ proteomics was performed that revealed 217 down-regulated and 177 up-regulated proteins in the combined Art and TMP group, indicating a significantly altered proteome profile as compared to that of the Art or TMP alone group. Functional enrichment analysis revealed the pharmacological effects of the combination of Art and TMP in the ECM mice brain proteome, such as axon growth, angiogenesis, and blood–brain transport. To the best of our knowledge, present study is the first comprehensive study to describe the brain proteomic alterations in ECM mice treated with Art, TMP and Art-TMP combination. This proteomic study not only provides the basis for further studies on mechanism of action of drugs, but also assists in identifying potential biomarkers for monitoring disease improvement of *Plasmodium* infection. At the same time, it enhances our understanding of the pathogenesis and host responses of this fatal parasitic disease. Further analysis involving patients after Art-TMP combination therapeutic interventions are required to provide additional insights into the correlation of the identified markers with the disease progression and their efficacy as disease monitoring or prognostic indicators, which could be an informative source for future persistence of the present investigation.

## Abbreviations

CM: cerebral malaria
Art: artesunate
TMP: Tetramethylpyrazine
pRBC: parasitized red blood cells
ECM: experimental cerebral malaria
SR: survival rate
RMCBS: rapid murine coma and behaviour scale
ICAM-1: intercellular adhesion molecule-1
VCAM-1: vascular cell adhesion molecule-1
GFAP: glial fibrillary acidic protein
NeuN: neuron-specific nuclear protein
BDNF: brain-derived neurotrophic factor
NT-3: neurotrophic factor-3
TNF-α: tumor necrosis factor
iTRAQ: isobaric tags for relative and absolute quantification
b-NGF: b-nerve growth factor
VEGF-A: vascular endothelial growth factor A

## Acknowledgement

No applicable

## Availability of data and materials

The datasets during and/or analysed during the current study available from the corresponding author on reasonable request.

## Authors’ contributions

YL, LC and XJ designed the research. XJ, LC, ZZ, YG, KL, TY, SQ and HL carried out experiments. XZ, LC, YC and XW provided guidance and access to materials and resources. YL, XJ and LC performed the analysis. YL, XJ and LC wrote the manuscript. All authors read and approved the final manuscript.

## Consent for publication

Not applicable.

## Author details

Artemisinin Research Center China Academy of Chinese Medical Sciences.

---

S1 Table. Differentially expressed proteins (DEPs) following treatment with artesunate (Art), tetramethylpyrazine (TMP), Art-TMP combination (Art+TMP).

S2 Table. Down- and up-regulated differentially expressed proteins (DEPs) in the artesunate (Art), tetramethylpyrazine (TMP) and Art-TMP combination (Art+TMP) groups.

S3 Table. The top 10 most significant brain differentially expressed proteins (DEPs) in the artesunate (Art), tetramethylpyrazine (TMP) and Art-TMP combination (Art+TMP) groups.

